# SIRT4 positively regulates autophagy *via* ULK1, but independently of HDAC6 and OPA1

**DOI:** 10.1101/2025.04.29.649368

**Authors:** Isabell Lehmkuhl, Khawar Amin, Nils Hampel, Afshin Iram, Julia Hesse, Constanze Wiek, Jasmin Thuy Vy Nguyen, Helmut Hanenberg, Jürgen Scheller, M. Reza Ahmadian, Björn Stork, Doreen M. Floss, Roland P. Piekorz

**Author notes:** Equal contribution (Isabell Lehmkuhl, Khawar Amin).

## Abstract

The sirtuin SIRT4 has been implicated in the control of autophagy and mitochondrial quality control *via* mitophagy, but the regulatory role of SIRT4 in autophagy/mitophagy induced by different stressors is unclear. Here, we show that cells expressing SIRT4(H161Y), a catalytically inactive, dominant-negative mutant of SIRT4, fail to upregulate LC3B-II and show a reduced autophagic flux upon treatment with different inducers of mitophagy/autophagy, *i.e*., CoCl_2_-triggered pseudohypoxia, CCCP/oligomycin-mediated respiratory chain inhibition, or rapamycin treatment. Interestingly, SIRT4(H161Y) expression (i) upregulated protein levels of HDAC6, which is involved in mitochondrial trafficking and autophagosome-lysosome fusion, and (ii) inhibited the conversion of OPA1-L to OPA1-S, which is associated with increased mitochondrial fusion and decreased mitophagy. Both HDAC6 and OPA1 are SIRT4 interactors. However, pharmacological inhibition of neither HDAC6 *via* Tubacin nor OPA1 *via* MYLS22 restored the stress-induced upregulation of LC3B-II levels upon autophagy/mitophagy treatment of SIRT4(H161Y)-expressing cells. Remarkably, inhibition of autophagosome-lysosome fusion and thus disruption of late autophagic flux by BafA1 treatment also failed to restore LC3B-II levels upon autophagy/mitophagy treatment, suggesting an inhibitory effect of SIRT4(H161Y) on the initiation/early phase of autophagy. Consistent with this idea, we show that SIRT4(H161Y) promoted the phosphorylation of ULK1 at S638 and S758 (mTORC1 targets), both of which mediate an important inhibitory regulation of autophagy initiation. Thus, our data suggest a positive regulatory function of SIRT4, presumably *via* modulation of AMPK/mTORC1 signaling, in the ULK1-dependent early regulation/initiation of stress-induced autophagic flux.

## Introduction

Autophagy is a critical cellular process that is triggered by a variety of extra/intracellular stresses, resulting in the degradation and recycling of cellular components. Among the latter are damaged organelles such as mitochondria, which are degraded *via* mitophagy to maintain mitochondrial homeostasis and function [1, 2]. A major inducer of autophagy is the rapamycin-mediated inhibition of the mammalian target of rapamycin complex 1 (mTORC1) [3], whereas increased mitophagy is triggered by hypoxia due to low oxygen levels or CoCl_2_-triggered pseudohypoxia [4]. Inhibition of the mitochondrial respiratory chain by treatment with carbonyl cyanide 3-chlorophenylhydrazone (CCCP) and oligomycin also results in a strong mitophagic response [5]. Dysregulation of autophagy and mitophagy plays an important role in several pathological conditions, including aging, cancer therapy resistance, cardiovascular diseases, and neurodegenerative disorders and therefore represents a common therapeutic target in these diseases [6].

Autophagy is a highly regulated process consisting of four phases, *i.e*., initiation controlled by the ULK1 initiation complex [7], elongation (phagophore formation and extension of autophagosomal membranes), maturation of autophagosomes, and finally degradation of contents following autophagosome-lysosome fusion [8, 9]. The elongation phase is associated with the microtubule-associated protein 1A/1B-light chain 3 (LC3) protein. LC3 is thereby widely used as a marker of autophagy due to its conversion from the cytosolic form (LC3-I) to the lipidated, autophagosomal membrane bound form (LC3-II) during autophagic flux [10]. At the regulatory level, autophagy is tightly controlled at its initiation phase by the Unc-51-like kinase 1 (ULK1) kinase [11]. Here, mTORC1 mediated phosphorylation of ULK at serine residues 758 and 638 disrupts AMPK-ULK1 interaction and therefore inhibits ULK1 dependent initiation of autophagy [12]. Thus, the mTORC1-ULK1 interaction controls initiation of autophagy.

Human sirtuins comprise a group of enzymes that mainly function in epigenetic regulation and gene expression control (SIRT1, SIRT2, SIRT6, and SIRT7; [13]), proliferation/cell survival and aging (*e.g.* SIRT6; [14]), and regulation of mitochondrial dynamics and metabolism (SIRT3, SIRT4, SIRT5; [15, 16]). SIRT4 is associated with cellular senescence and mitophagy [17, 18]. At the molecular level, SIRT4 interacts with the mitochondrial GTPase OPA1 and stabilizes its inner-membrane bound long form (OPA1-L) [17]. In contrast to the short form OPA1-S, OPA1-L promotes mitochondrial fusion and thereby counteracts fission that is typically required for mitophagy [19]. At the same time, SIRT4 inhibits MAPK mediated phosphorylation and activation of DRP1 leading to impaired mitochondrial fission [20]. The molecular interactions of SIRT4 with DRP1 and OPA1 are not well understood, and it is unknown, to which extent they are linked to impair mitophagy caused by (pseudo)hypoxia or inhibition of mitochondrial chain function. Another recently described SIRT4 interactor, histone deacetylase 6 (HDAC6) [21, 22], is known to regulate autophagy in terms of autophagosome transport and autophagosome-lysosome fusion [23, 24]. However, also here it is unclear whether and how the SIRT4-HDAC6 interaction plays a role in regulation of autophagy/mitophagy triggered by different stressors.

The present study employs various autophagy and mitophagy stressors to define the role of SIRT4 in the regulation of autophagy/mitophagy. We observed an overall inhibitory impact of SIRT4(H161Y), an enzymatically inactive, dominant-negative SIRT4 mutant, on LC3B-II upregulation and autophagy. This phenotype could not be rescued by pharmacological inhibition of OPA1-L and HDAC6, that were both upregulated upon SIRT4(H161Y) expression. However, in contrast to wild-type SIRT4, SIRT4(H161Y) increased inhibitory phosphorylation of ULK1. Thus, our results suggest that SIRT4 has a positive effect on ULK1 and early autophagy regulation, presumably through the AMPK-mTORC1-ULK1 network.

## Materials and Methods

### Reagents and antibodies

The following reagents were obtained from the indicated companies: CoCl_2_ (Roth), Bafilomycin A_1_ (BafA_1_) and tubacin (Cayman Chemical), CCCP (carbonyl cyanide 3-chlorophenylhydrazone; Sigma-Aldrich), oligomycin and rapamycin (Biomol and Tocris), and MYLS22 (MedChemExpress EU). Primary antibodies used in immunoblot analysis are listed in **Table S1**. Anti-mouse (700 nm; LI-COR IRDye #926-32213) or anti-rabbit (800 nm; LI-COR IRDye #926-68072) were used as secondary antibodies.

### Cell culture

Parental and SIRT4 wild-type/mutant expressing HEK293 cell lines were cultured at 37°C and 5% CO_2_ in DMEM (Dulbecco’s Modified Eagle Medium) containing high glucose (4.5 g/L; Thermo Fisher Scientific) with 10% fetal bovine serum (FBS; Gibco) and penicillin (100 units/mL)/streptomycin (100 μg/mL) (Genaxxon). HEK293 cells were obtained from the German Collection of Microorganisms and Cell Cultures GmbH (DSMZ) (HEK293: ACC 305). HEK293-eGFP, HEK293-SIRT4-eGFP, and HEK293-SIRT4(H161Y)-eGFP cell lines as well as HEK293 cell lines stably expressing Myc-Flag, SIRT4-Myc-Flag, or SIRT4(H161Y)-Myc-Flag have been described previously [17, 21] and were grown under selection with 400 µg/ml Geneticin/G418 (Genaxxon)

### Treatment of HEK293 cell lines with mitophagy or autophagy inducers and selected inhibitors

HEK293 cell lines were subjected to mitophagy induction in a chemical hypoxia/pseudohypoxia model [4, 25] using CoCl_2_ treatment at a concentration of 400 µM for 24 h or as indicated. Alternatively, mitophagy was triggered by mitochondrial membrane depolarization upon combined treatment with CCCP (5 µM) and oligomycin (5 µM) for 24 h. General autophagy was induced by rapamycin treatment (1.25 µM, 24 h). Inhibitors against HDAC6 (tubacin) and OPA1 (MYLS22) were used for a (co)treatment period of 24 h at concentrations of 2 µM and 50 µM, respectively. BafA_1_, an inhibitor of autophagosome-lysosome fusion, was used for a (co)treatment period of 4 h at a concentration of 0.5 µM.

### Preparation of total cell lysates and immunoblot analysis

Total cell lysates were generated using lysis buffer containing 0.3% CHAPS (3-[(3-Cholamidopropyl) dimethylammonio]-1-propanesulfonate), 50 mM Tris-HCl (pH 7.4), 150 mM NaCl, 1 mM Na_3_VO_4_, 10 mM NaF, 1 mM EDTA, 1 mM EGTA, 2.5 mM Na_4_O_7_P_2_, and 1 µM DTT. The cOmplete™ protease inhibitor cocktail (Sigma-Aldrich) was used to prevent the degradation of proteins in the lysates. The latter were cleared by centrifugation (11.000 x g at 4°C for 20 min) and the protein concentration of the supernatants (total cell lysates) was determined using the Bradford assay (Roth). Relative quantification of protein levels (as compared to α-tubulin or β-actin protein loading controls) was performed by ImageJ-based densitometric analysis of specific immunoblot signals.

### Flow cytometry-based measurement of autophagic flux using the GFP-LC3-RFP-LC3ΔG probe

The expression vector pMRX-IP-GFP-LC3-RFP-LC3(ΔG) (addgene plasmid #84572) [26]) was transfected into HEK293 cell lines stably expressing Myc-Flag, SIRT4-Myc-Flag, or SIRT4(H161Y)-Myc-Flag [21] using the Turbofect transfection reagent (Thermo Fisher Scientific). Stable cell lines expressing GFP-LC3-RFP-LC3(ΔG) were grown in double selection media (400 µg/ml geneticin/G418; Genaxxon; 1.5 µg/ml puromycin; InvivoGen). GFP-LC3-RFP-LC3ΔG works as an autophagic flux probe (GFP-LC3) that upon ATG4-mediated cleavage releases both GFP-LC3 and RFP-LC3ΔG. In contrast to GFP-LC3, RFP-LC3ΔG cannot be incorporated into the autophagosomal membrane and therefore serves as cytosolic reference control [26]. The experimental GFP/RFP ratios, which decrease upon induction of autophagic flow that is followed by LC3B-II degradation upon autophagosome-lysosome fusion, were measured by flow cytometry (BD FACS Canto II). Results were analyzed using FlowJo v10 software and calculated as MFI (Median Fluorescence Intensity).

### Flow cytometry-based measurement of mitophagy using the pH-sensitive and mitochondrially (mt) targeted mt-mKEIMA probe

The retroviral plasmid pCHAC-mt-mKEIMA (addgene plasmid #72342) [27]) was stably transduced into HEK293 cell lines stably expressing eGFP, SIRT4-eGFP, or SIRT4(H161Y)-eGFP [21] essentially as described [28]. Upon autophagosome-lysosome fusion and hence low auto-lysosomal pH, mt-mKEIMA (excitation/emission maxima λ=586/620 nm at a pH < 5) can be as employed as quantitative marker of mitophagic activity [29]. Analysis was performed by flow cytometry (BD LSRFortessa) and results were evaluated using the FlowJo v10 software and calculated as MFI (Median Fluorescence Intensity).

### Cellular analysis of mitochondrial mass using the MitoMark Green I probe

Mitochondrial content was determined by flow cytometry (BD FACS Canto II) using MitoMark Green I staining (biotechne-Tocris; excitation/emission maxima λ ∼ 490/516 nm). MitoMark Green I localizes to mitochondria independent of their membrane potential. Results were analyzed using the FlowJo v10 software and calculated as MFI (Median Fluorescence Intensity).

### Statistical analysis

Data are presented as mean ± S.D. Multiple comparisons were analyzed by one-way or two-way analysis of variance (ANOVA) followed by Tukey’s post-hoc test to identify group differences in variance analysis using the GraphPad Prism software. Statistical significance was set at the level of p ≤ 0.05 (*p ≤ 0.05, **p ≤ 0.01, and ***p ≤ 0.001).

## Results

### The enzymatically inactive SIRT4 mutant SIRT4(H161Y) inhibits stress-induced LC3B-II generation, autophagic flux, and mitophagy

First of all, we performed a comparative characterization of the role of SIRT4 in the regulation of mitophagy/autophagy caused by different stressors, in particular CoCl_2_-induced pseudohypoxia that activates HIF1α and upregulates LC3B-II (**Fig. S1**). We employed HEK293 cells stably expressing eGFP (vector control), wild-type SIRT4-eGFP, or its catalytically inactive mutant SIRT4(H161Y)-eGFP (**Fig. S2A, B**) [17, 25]. Interestingly, among the cell lines analyzed, HEK293-SIRT4(HY)-eGFP cells showed strongly reduced LC3B-II levels upon CoCl_2_ treatment for 24 h (**Fig. 1A, B**), whereas the levels of LCB-I were comparable between the genotypes analyzed (**Fig. 1C**). Similar results were obtained for LC3B-II when treating these cell lines with CoCl_2_ for 36 h (**Fig. S3**). We extended these findings to flow cytometry-based measurement of autophagic flux using the GFP-LC3-RFP-LC3ΔG reporter probe [26], for which HEK293 cells stably expressing SIRT4-myc-Flag or its catalytically inactive mutant SIRT4(H161Y)-myc-Flag were generated (**Fig. S2C, D**). The results of these experiments were consistent with the LC3B-II immunoblot analyses, observing the weakest CoCl_2_-triggered reduction in the GFP-LC3/RFP-LC3ΔG ratio (*i.e*., the lowest autophagic flux induction) for SIRT4(H161Y)-myc-Flag expressing cells (**Fig. 1D**). Moreover, flow cytometric analyses using mitochondria specific readouts showed that the CoCl_2_ triggered increase in mitophagic activity (mt-mKEIMA probe) was lowest in SIRT4(H161Y) expressing cells, concomitant with the strongest relative increase in mitochondrial content (Mitomark Green I probe) (**Fig. 1E, F**). Of note, the SIRT4(H161Y) mediated inhibition of autophagic flux was also observed in GFP-LC3/RFP-LC3ΔG ratio measurements upon CCCP/oligomycin triggered inhibition of mitochondrial membrane potential, resulting in mitophagy, as well as upon general autophagy induced by rapamycin-mediated mTORC1 inhibition (**Fig. S4A, B**). The latter was confirmed through effective down-regulation of pS6, a target of the mTORC1 – p70S6-kinase (p70S6K) axis (**Fig. S4C**). Taken together, expression of the dominant-negative SIRT4(H161Y), and this at lower protein levels as compared to ectopic wild-type SIRT4 (**Fig. S2A, C**), impairs autophagic flux upon CoCl_2_-induced pseudohypoxia, mitochondrial membrane depolarization, or rapamycin treatment.

**Fig. 1.**
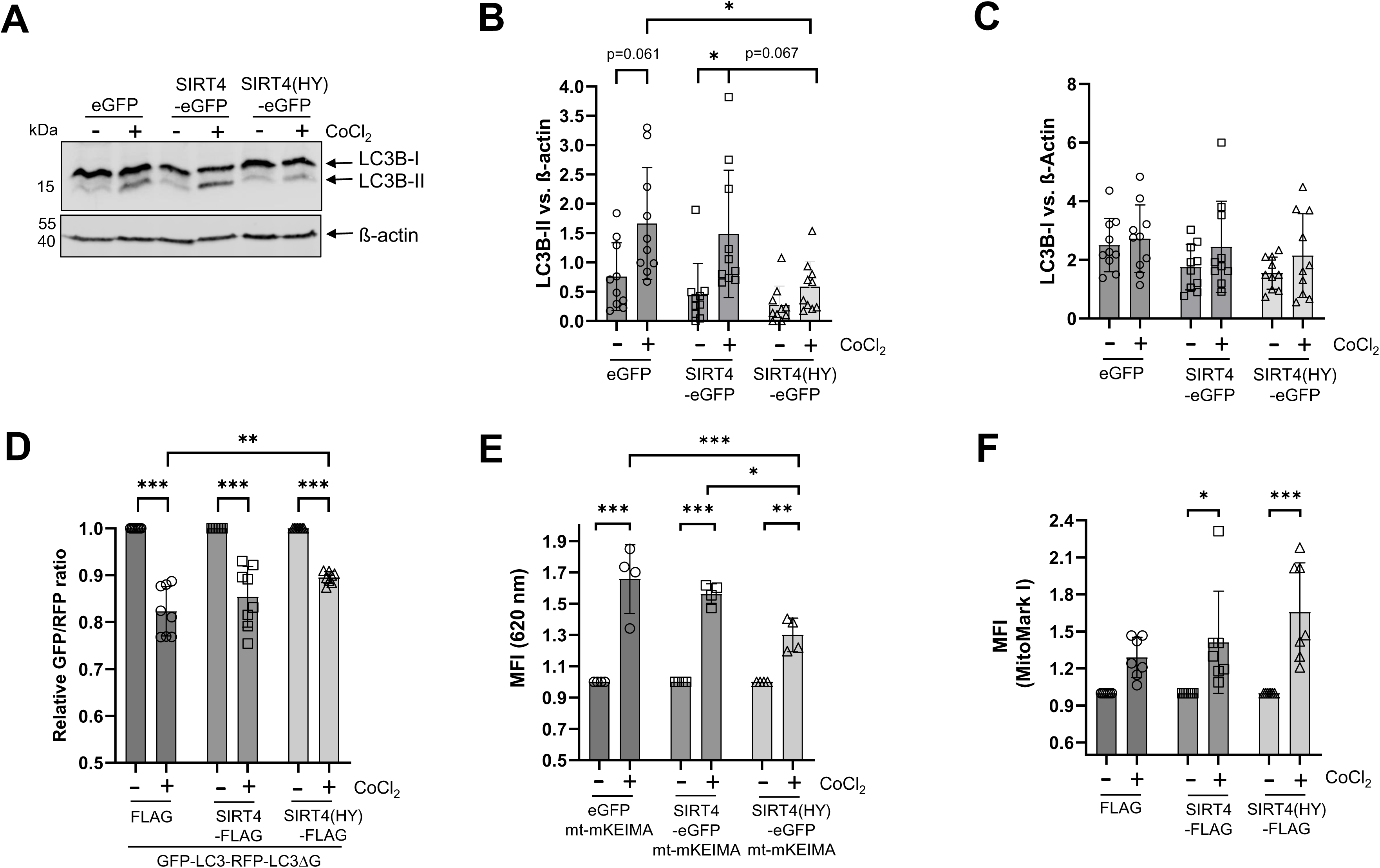
Inhibitory impact of SIRT4(H161Y) on autophagic flux and mitophagy upon CoCl_2_-induced pseudohypoxia. (**A**) HEK293 cells stably expressing eGFP, SIRT4-eGFP, or SIRT4(H161Y)-eGFP were subjected to CoCl_2_-induced hypoxia followed by immunoblot analysis of LC3B-I/II levels (n=10). (**B, C**) Relative quantification of immunoblot signals of LC3B-II and LC3B-I was performed using ImageJ-based densitometric evaluation and β-actin levels as loading control. (**D**) HEK293 cells stably expressing myc-Flag, SIRT4-myc-Flag, or SIRT4(H161Y)-myc-Flag were subjected to CoCl_2_-induced hypoxia followed by flow cytometry-based analysis of autophagic flux using the GFP-LC3-RFP-LC3ΔG probe (n=8). The more the GFP-LC3/RFP-LC3ΔG ratio decreases, the higher is the autophagic flux. (**E**) Analysis of mitophagic activity using the mt-mKEIMA fluorescent protein (n=4). (**F**) Mitochondrial content analysis using the MitoMark Green I probe (n=7). To determine statistical significance, Two-Way ANOVA tests were employed (mean ± S.D.; *p < 0.05; **p < 0.01; ***p < 0.001). MFI: Median fluorescent intensity.

### SIRT4(H161Y)-mediated inhibition of CoCl_2_-induced LC3B-II increase and autophagic flux is not linked to increased HDAC6 levels and decreased acetylated α-Tubulin (acK40)

The autophagy regulator HDAC6 interacts with SIRT4 [21, 22] and is also found at increased levels upon expression of SIRT4 or SIRT4(H161Y) (**Fig. 2A, B**). In contrast to eGFP expressing controls, SIRT4 and SIRT4(HY) expressing cells maintained high HDAC6 protein levels and did not downregulate HDAC6 upon CoCl_2_ treatment (**Fig. 2C**). Concomitantly, up-regulated HDAC6 caused down-regulation of acetylated α-Tubulin (acK40), which is a major HDAC6 target (**Fig. 2D, E**). Acetylated α-tubulin (acK40) is required for efficient microtubule dynamics and transport processes in connection with autophagy/mitophagy. Given that pharmacological inhibition of HDAC6 can restore autophagy in various experimental model systems [30–33], we next addressed the effect of the HDAC6 inhibitor tubacin on autophagic flux in SIRT4(H161Y) expressing cells. As expected, tubacin treatment resulted in clear upregulation of acetylated α-tubulin (K40) levels in all genotypes analyzed (**Fig. S5**). However, at the same time, inhibition of HDAC6 failed to rescue CoCl_2_-induced LC3B-II increase, autophagic flux, and mitophagy in HEK293-SIRT4(H161Y) cells (**Fig. 3**). Of note, co-treatment with tubacin & CCCP/oligomycin (**Fig. S6A**), but not tubacin and rapamycin (**Fig. S7A**), revealed a slight and significant rescue in autophagic flux as measured through the flow cytometric GFP-LC3/RFP-LC3ΔG ratio method. We conclude that the increased HDAC6 levels are not decisive for the impaired LC3B-II/autophagic flux in SIRT4/SIRT4(H161Y) expressing HEK293 cells upon induction of mitophagy or autophagy.

**Fig. 2:**
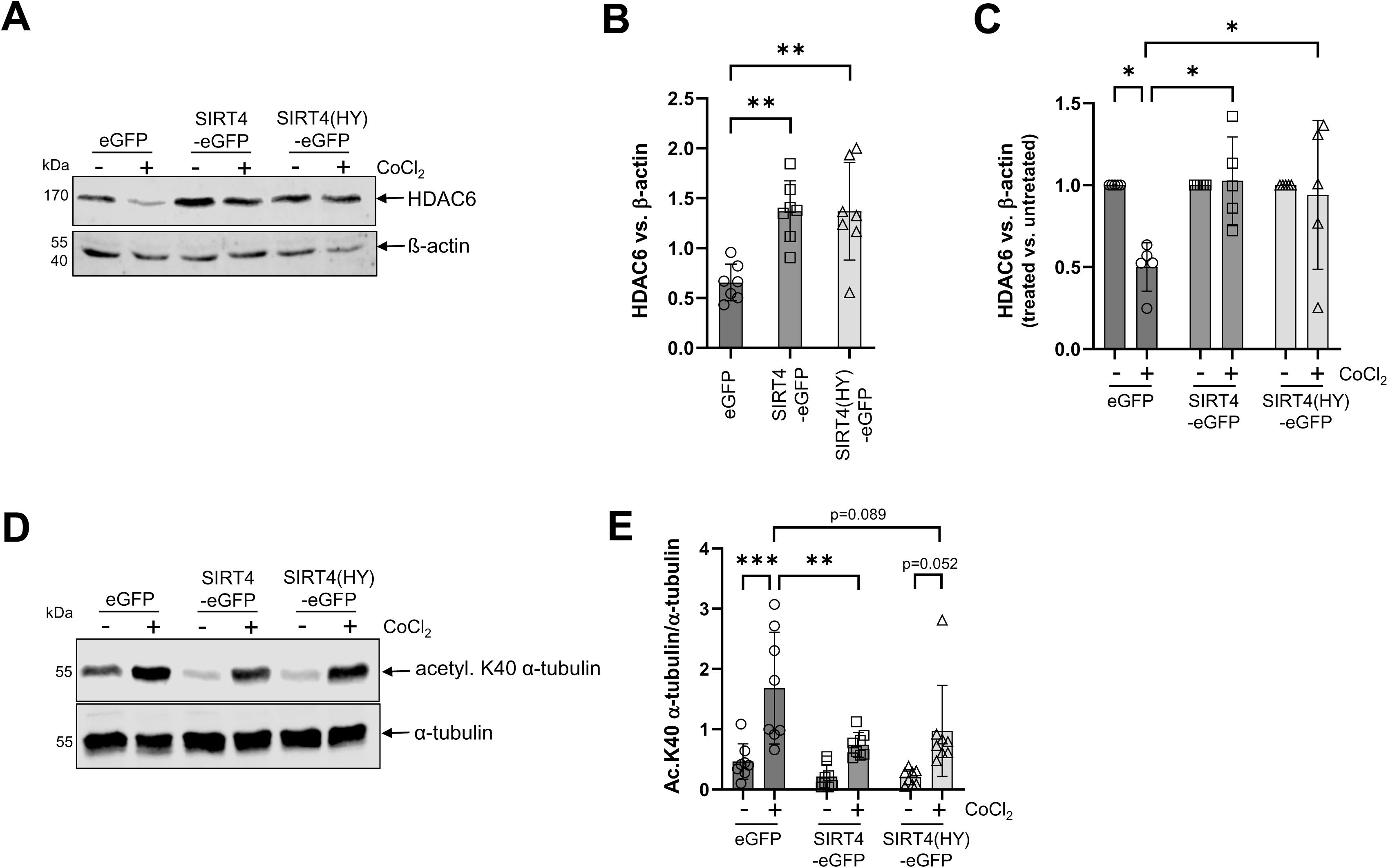
Ectopic expression of SIRT4 or SIRT4(H161Y) leads to upregulation of HDCA6 protein levels and concomitant reduced acetylated α-tubulin (K40). (**A**) Immunoblot and densitometric (**B**, **C**) analysis of HDAC6 protein levels in untreated and CoCl_2_-treated HEK293 cells expressing SIRT4 or SIRT4(H161Y) (n=5 to 7). Immunoblot (**D**) and densitometric (**E**) analysis of acetylated α-tubulin (K40) protein levels in untreated and CoCl_2_-treated HEK293 cells expressing SIRT4 or SIRT4(H161Y) (n=8). To determine statistical significance, Two-Way ANOVA tests were employed (mean ± S.D.; *p < 0.05; **p < 0.01; ***p < 0.001).

**Fig. 3.**
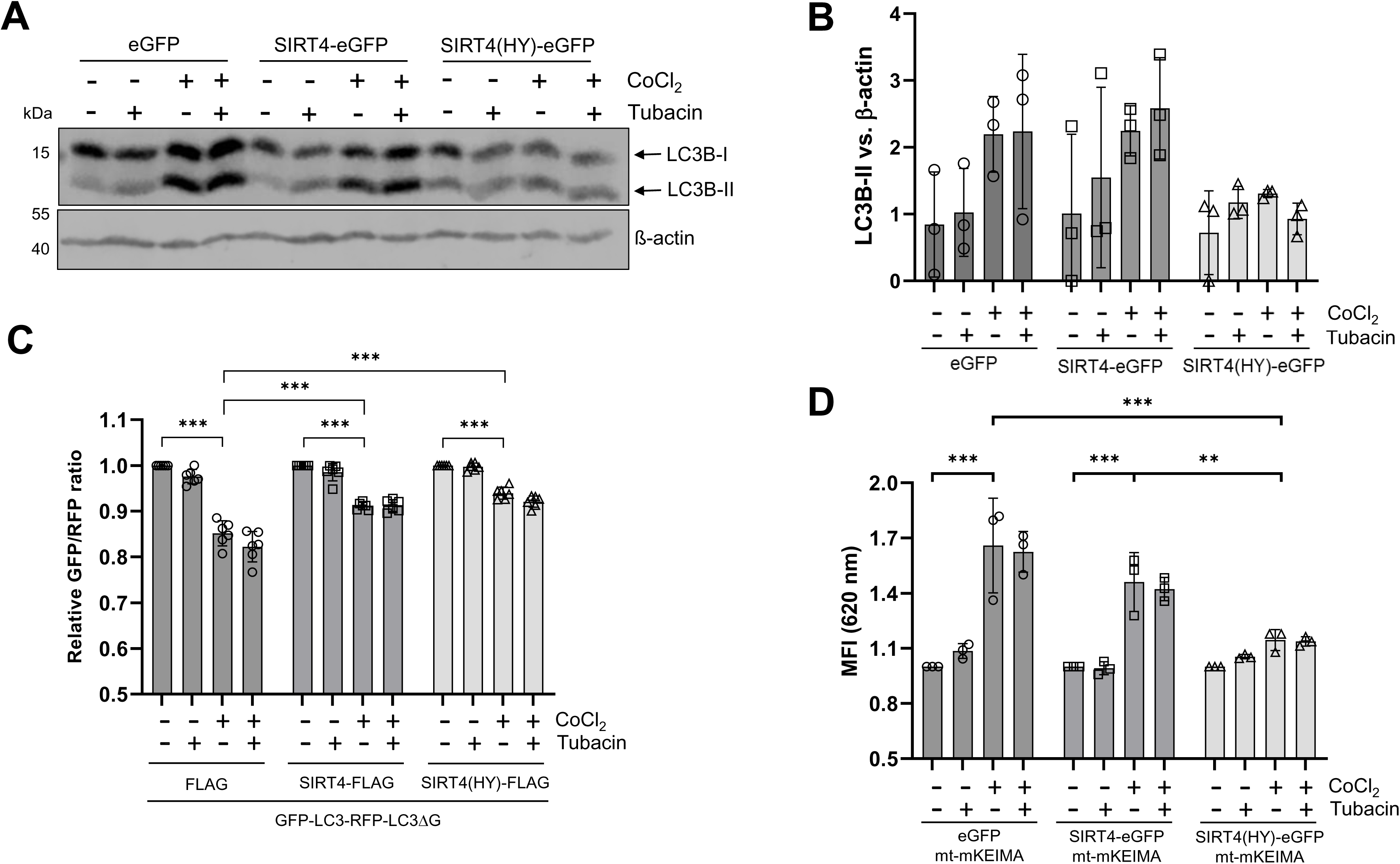
The HDAC6 inhibitor tubacin fails to rescue LC3B-II levels, autophagic flux, and mitophagy in SIRT4(H161Y) expressing HEK293 cells upon CoCl_2_-induced pseudohypoxia. (**A**) HEK293 cells stably expressing eGFP, SIRT4-eGFP, or SIRT4(H161Y)-eGFP were subjected to CoCl_2_-induced hypoxia in the presence or absence of the HDAC6 inhibitor Tubacin followed by immunoblot analysis of LC3B-I/II levels. (**B**) Relative quantification of immunoblot signals of LC3B-II was performed using ImageJ-based densitometric evaluation and β-actin levels as loading control (n=3). (**C**, **D**) HEK293 cells stably expressing FLAG, SIRT4-myc-Flag, or SIRT4(H161Y)-myc-Flag were subjected to CoCl_2_-induced hypoxia in the presence or absence of tubacin followed by flow cytometry-based analysis of autophagic flux (**C**; n=6) and mitophagic activity (**D**; n=3). To determine statistical significance, Two-Way ANOVA tests were employed (mean ± S.D.; **p < 0.01; *** p < 0.001). MFI: Median fluorescent intensity.

### Lack of CoCl_2_-induced LC3B-II increase and autophagic flux in SIRT4(H161Y) expressing cells is not based on increased OPA1-L stabilization and altered mitochondrial dynamics

SIRT4 interacts with the mitochondrial GTPase OPA1 and stabilizes its inner-membrane bound long form (OPA1-L) [17] that promotes mitochondrial fusion and thereby potentially counteracts mitophagy as compared to OPA1-S [19, 34]. Upon CoCl_2_ treatment, HEK293 cells expressing SIRT4(H161Y) maintained the highest OPA1-L levels and therefore OPA1-L/OPA1-S ratios as analyzed over a period of 36 h (**Fig. 4A, B**). Next, we tested if the increased OPA1-L may affect LC3B-II upregulation. Therefore, HEK293 SIRT4(H161Y) cells were subjected to co-treatment with the OPA1 inhibitor MYLS22 [35]. As indicated in **Fig. 4C, D**, MYLS22 reduced the OPA1-L/OPA1-S ratio in SIRT4(H161Y) expressing cells, although there was no additive/synergistic effect with CoCl_2_ co-treatment. However, although MYLS22 alone did up-regulate LC3B-II levels, co-treatment using MYLS22 and CoCl_2_ failed to significantly rescue LC3B-II (**Fig. 5A, B**), autophagic flux, and mitophagy (**Fig. 5C, D**) in CoCl_2_ treated HEK293-SIRT4(HY) cells. In addition, co-treatment using MYLS22 and CCCP/oligomycin (**Fig. S6B**) or MYLS22 and rapamycin (**Fig. S7B**) did not significantly increase autophagic flux measured by the GFP-LC3/RFP-LC3ΔG ratio-based method. Thus, our data indicate that altered mitochondrial dynamics is not the reason for the impaired LC3B-II/autophagic flux in SIRT4(H161Y) expressing HEK293 cells upon induction of mitophagy or autophagy.

**Fig. 4.**
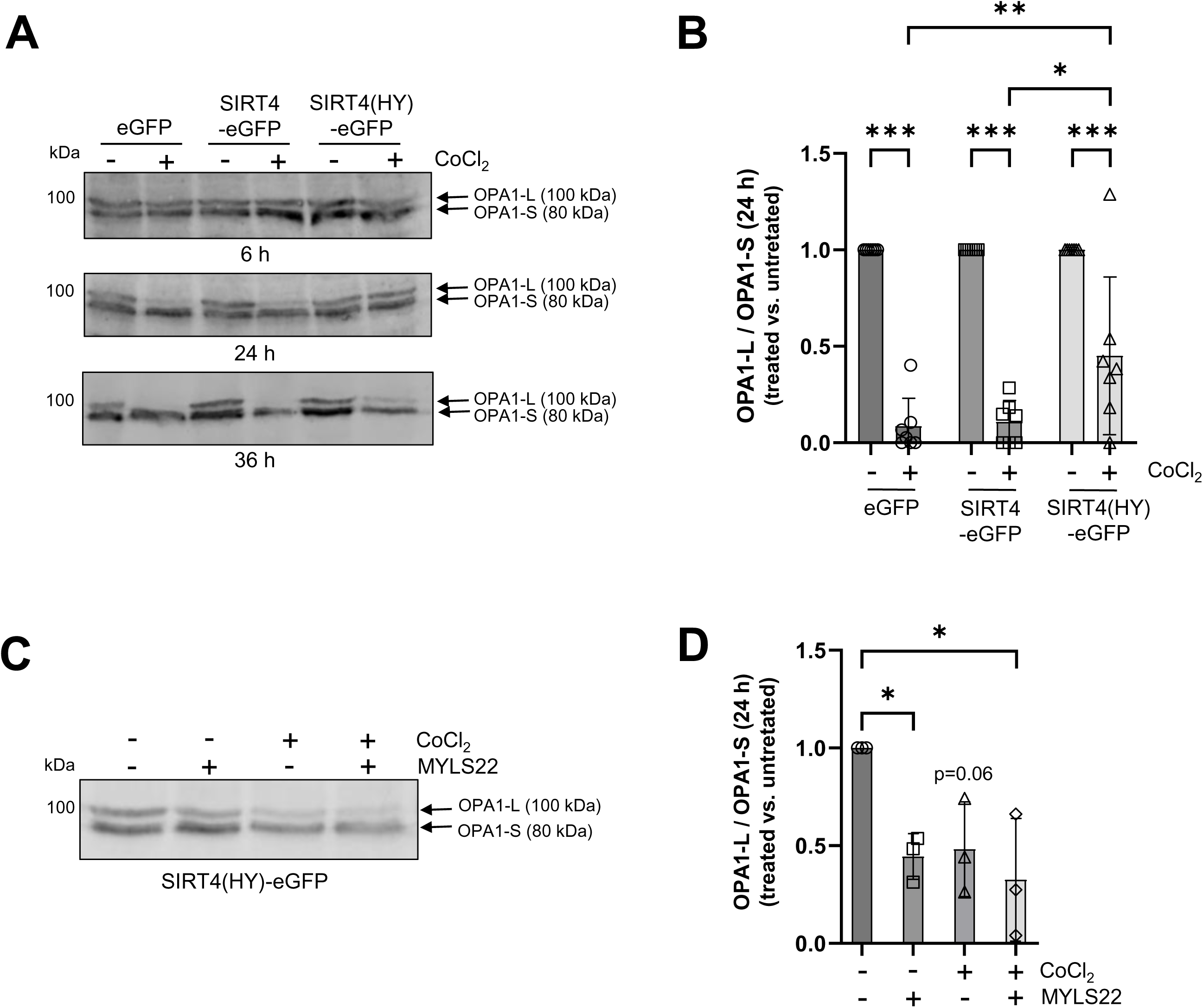
SIRT4(H161Y) inhibits CoCl_2_-induced conversion of OPA1-L to OPA1-S. (**A**) Immunoblot and densitometric analysis (**B**) of OPA1-L/OPA1-S protein levels in untreated and CoCl_2_-treated HEK293 cells expressing SIRT4 or SIRT4(H161Y) (n=6). (**C**) The OPA1 inhibitor MYLS22 decreases the OPA1-L/OPA1-S protein ratio as quantified by densitometric analysis (**D**; n=3). Cell treatment was performed for 24 hrs. To determine statistical significance, Two-Way ANOVA tests were employed (mean ± S.D.; * *p* < 0.05; ** *p* < 0.01; *** p < 0.001).

**Fig. 5.**
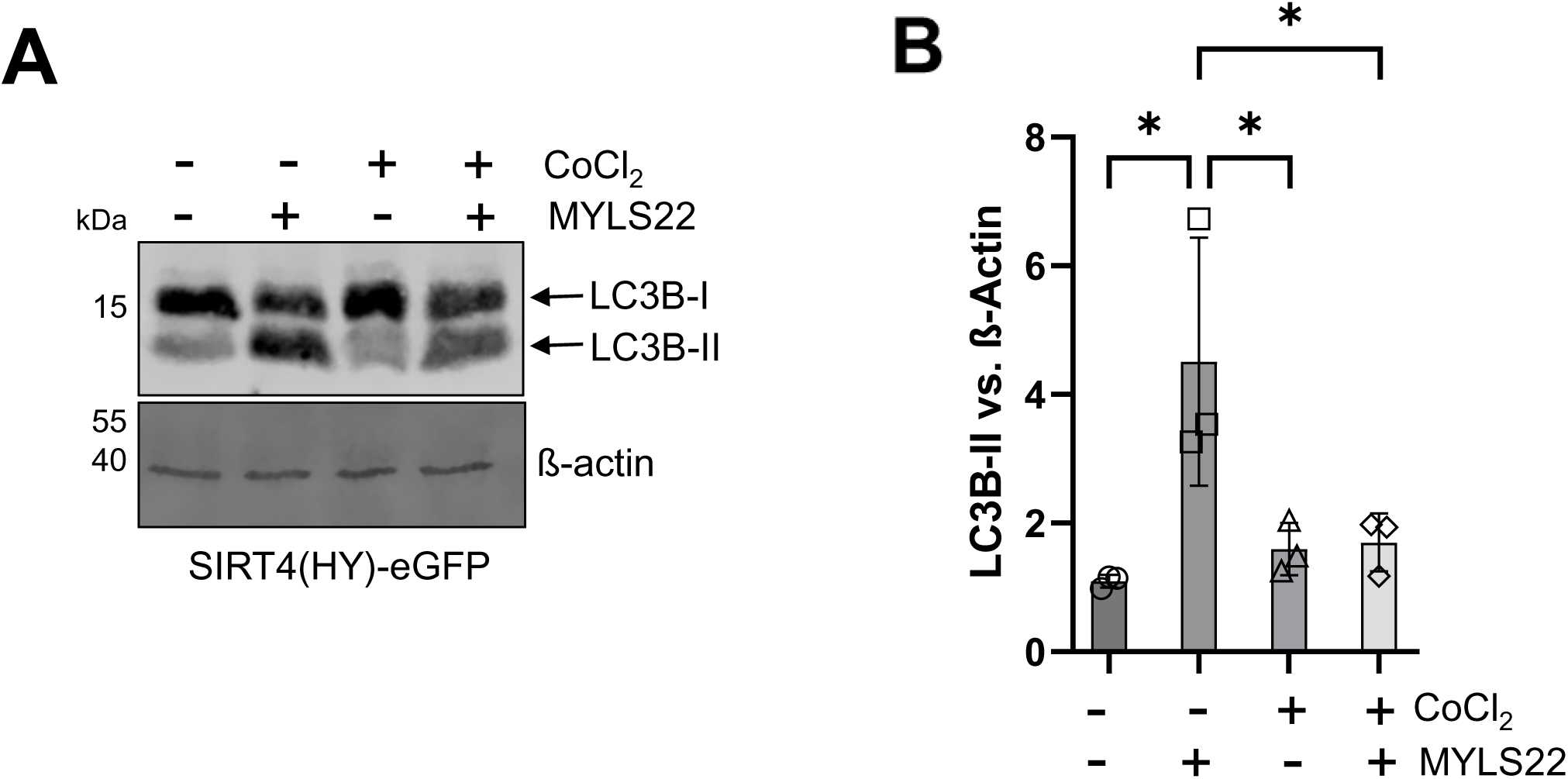
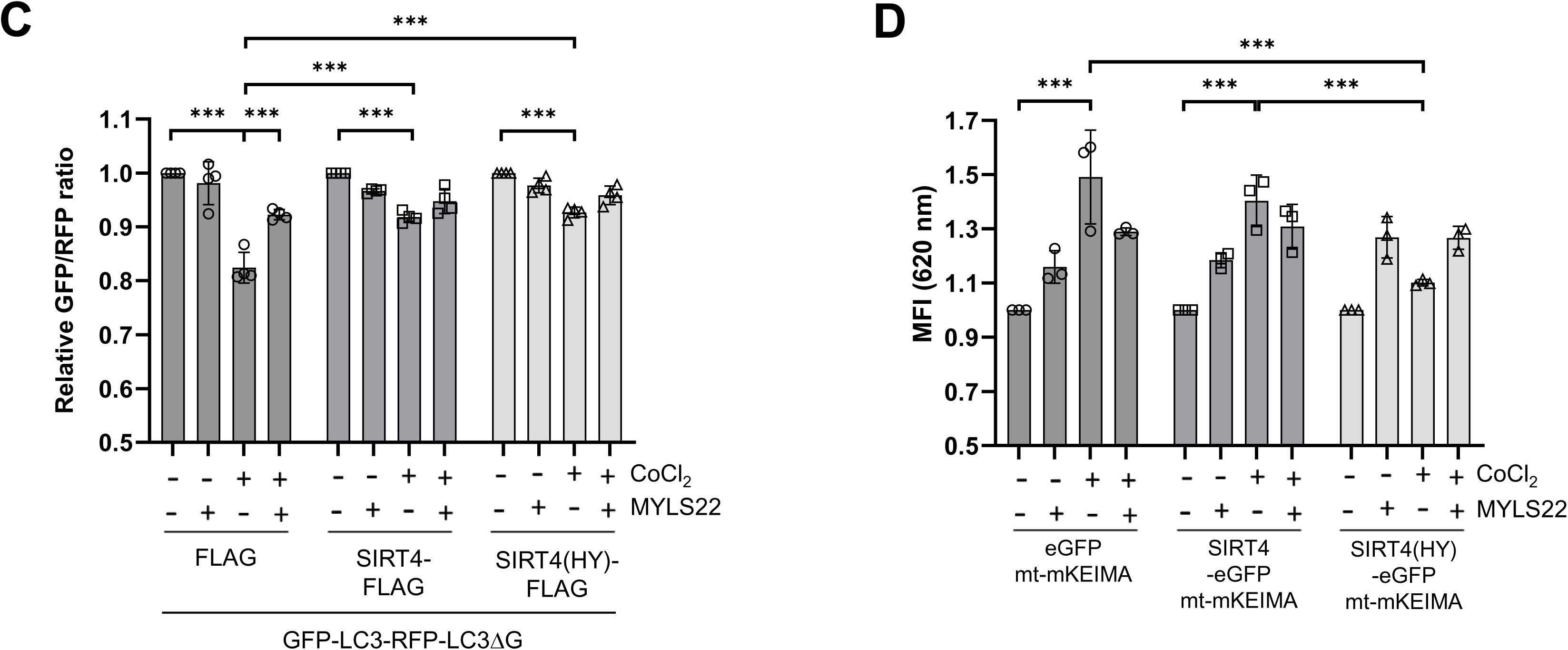
The OPA1 inhibitor MYLS22 fails to restore LC3B-II levels, autophagic flux, and mitophagy in SIRT4(H161Y) expressing HEK293 cells upon CoCl_2_-induced pseudohypoxia. (**A**) Immunoblot and (**B**) densitometric analysis of LC3B-II protein levels in untreated and CoCl_2_-treated HEK293 cells expressing SIRT4 or SIRT4(H161Y) in the presence or absence of MYLS22 (n=3). (**C**) HEK293 cells stably expressing myc-Flag, SIRT4-myc-Flag, or SIRT4(H161Y)-myc-Flag were subjected to CoCl_2_-induced hypoxia in the presence or absence of MYLS22 followed by flow cytometry-based analysis of autophagic flux using the GFP-LC3-RFP-LC3ΔG probe (n=4). (**D**) HEK293 cells stably expressing eGFP, SIRT4-eGFP, or SIRT4(H161Y)-eGFP were subjected to CoCl_2_-induced hypoxia in the presence or absence of MYLS22 followed by flow cytometry-based analysis of mitophagy using the mt-mKEIMA probe (n=3). To determine statistical significance, One-Way or Two-Way ANOVA tests were employed (mean ± S.D.; *p < 0.05; ***p < 0.001). MFI: Median fluorescent intensity.

### BafA_1_-mediated inhibition of autophagosome-lysosome fusion fails to restore LC3B-II levels in CoCl_2_-treated SIRT4(H161Y) expressing cells

Execution of autophagy depends on fusion of autophagosomes with lysosomes that can be inhibited by the V-ATPase and SERCA inhibitor BafA_1_ [36–38]. BafA_1_ treatment is widely used to address the basis for increased LC3B-II levels and autophagic flux, *i.e*. to differentiate between increased biosynthesis/modification of LC3B-I to LC3B-II *vs.* increased auto-lysosomal degradation of LC3B-II [38]. Interestingly, under basal conditions, SIRT4(H161Y) expressing HEK293 cells displayed an approximately three-fold increase in LC3B-II upon BafA_1_ treatment for a period of 4 h (**Fig. 6**). However, upon combined CoCl_2_-treatment, BafA_1_ failed to significantly increase LC3B-II levels (**Fig. 6**), indicating that an accelerated degradation of LC3B-II by the autolysosome does not account for the observed phenotype. These results suggest that the stress-induced generation of LCB-II is inhibited upon SIRT4(H161Y) expression at an earlier stage, *i.e.,* at the LC3B-I to LC3BII conversion or even at the initiation of autophagy.

**Fig. 6.**
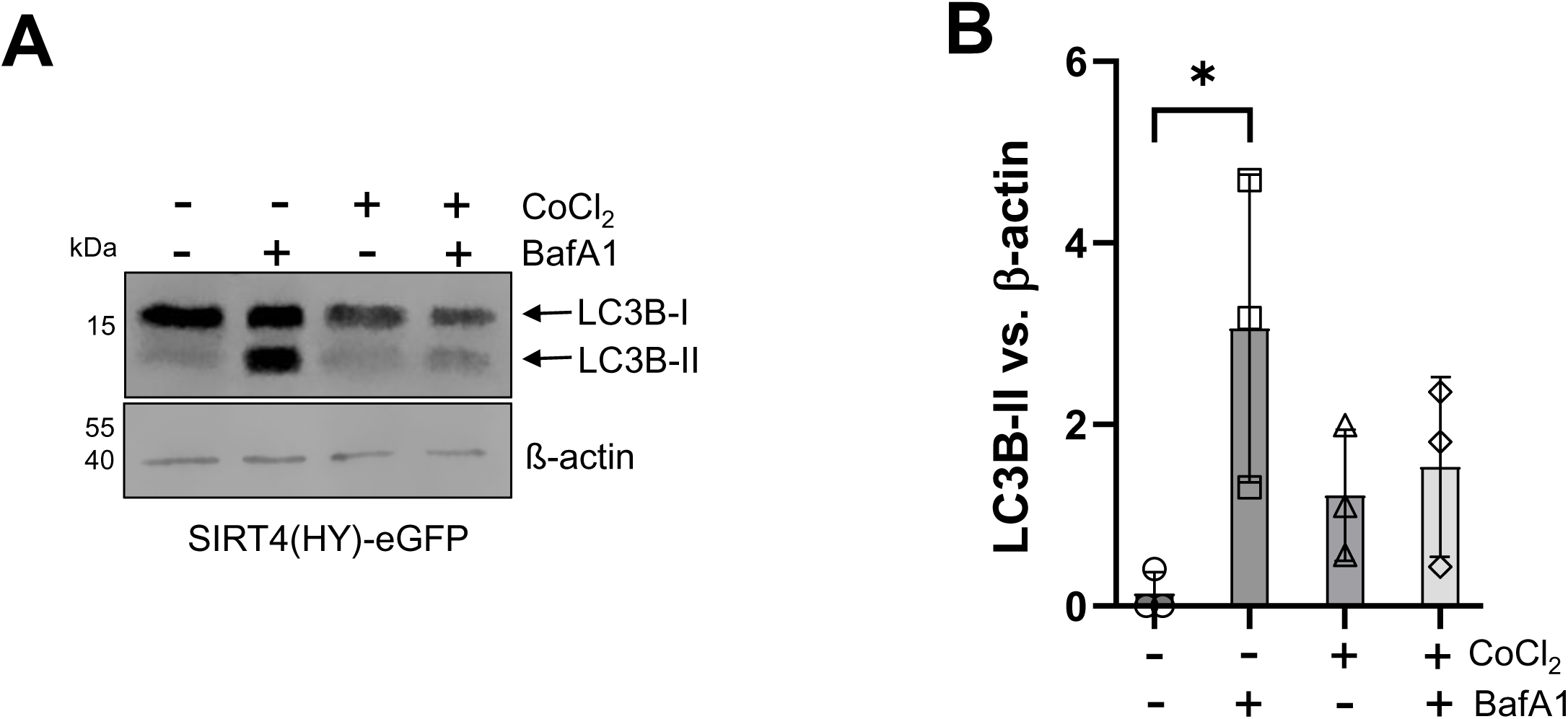
BafA_1_ treatment does not restore LC3II-B levels in SIRT4(H161Y) expressing HEK293 cells upon CoCl_2_-induced pseudohypoxia. (**A**) Immunoblot analysis and (**B**) densitometric evaluation of LC3B-II *vs*. β-actin as loading control was performed (n=3). To determine statistical significance, One-Way ANOVA tests were employed (mean ± S.D.; *p < 0.05).

### SIRT4(H161Y) promotes phosphorylation of ULK-1 at Ser757 and Ser638 that are linked to negative regulation of early autophagy

Major upstream regulators of autophagy include mTORC1 that affects the central ULK1-driven autophagy initiation complex in a negative manner [7–9, 11, 39, 40]. Given the interaction of SIRT4 with the AMPK/mTORC1 pathway [41–43], we asked whether the early regulatory phase of autophagy may be affected by ectopic expression of the dominant-negative mutant SIRT4(H161Y). Therefore, we analyzed the relative levels of ULK1 phosphorylated at serine residues Ser758 and Ser638 that both represent mTORC1 substrates [7, 12, 44], associated with inhibition of ULK1 function and hence decreased initiation of autophagy. As indicated in **Figure 7**, combined immunoblotting and densitometric analysis revealed that CoCl_2_ treatment resulted in downregulation of ULK1^pS758^ and ULK^pS638^ levels in HEK293 cells that express eGFP or SIRT4-eGFP. In contrast, in SIRT4(H161Y)-GFP expressing HEK293 cells, CoCl_2_ failed to reduce the amount of both ULK1^pS758^ and ULK^pS638^ as compared to the untreated controls. In conclusion, our findings suggest that expression of SIRT4(H161Y) negatively interferes with ULK1 dephosphorylation in response to stressors that induce autophagy.

**Fig. 7.**
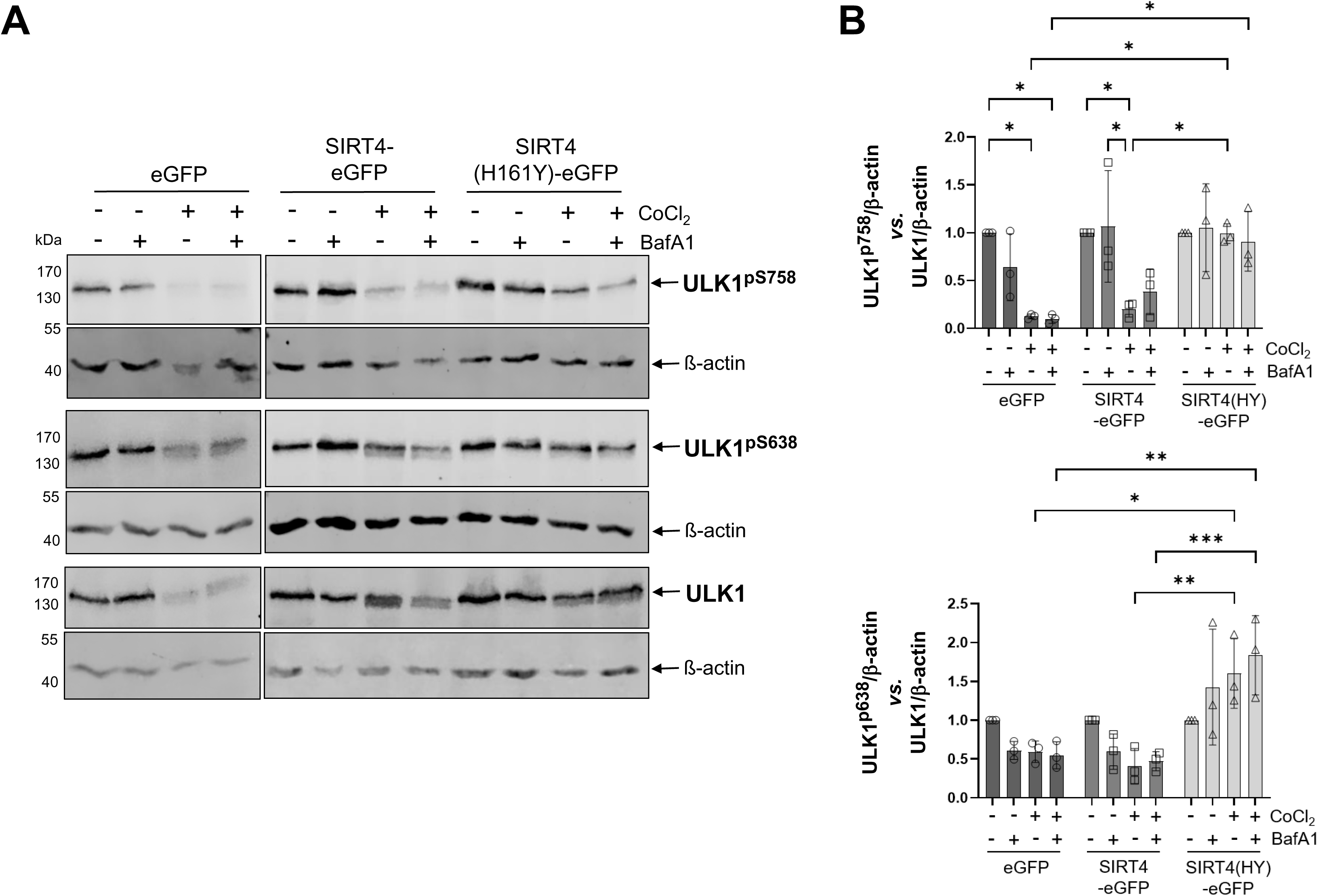
SIRT4(H161Y) increases inhibitory phosphorylation of ULK1 upon CoCl_2_-induced pseudohypoxia. (**A**) HEK293 cells stably expressing eGFP, SIRT4-eGFP, or SIRT4(H161Y)-eGFP were subjected to CoCl_2_-induced hypoxia followed by immunoblot analysis of ULK1^pS758^ and ULK1^pS638^ levels. (**B**) Relative quantification of immunoblot signals of ULK1^pS758^ (upper panel) and ULK1^pS638^ (lower panel) was performed using ImageJ-based densitometric evaluation and β-actin levels as loading control (n=3). To determine statistical significance, Two-Way ANOVA tests were employed (mean ± S.D.; * *p* < 0.05; ** *p* < 0.01; *** p < 0.001).

## Discussion

This study provides novel insight into the role of the sirtuin SIRT4 in the regulation of autophagy/mitophagy. In summary, we showed that (i) the dominant-negative mutant SIRT4(H161Y)-eGFP prevents LC3B-II upregulation upon treatment with various inducers of mitophagy/autophagy; (ii) mechanistically, this phenotype is independent of the SIRT4 interactors OPA1 and HDAC6 as shown by pharmacological inhibition experiments; (iii) the low LC3B-II levels due to SIRT4(H161Y) expression could not be restored by BafA_1_-mediated inhibition of autophagosome-lysosome fusion, therefore ruling out an increased auto-lysosomal degradation of LC3B-II at the late phase of autophagy; (iv) interestingly, expression of SIRT4(H161Y) increased the levels of UKL1 kinase phosphorylated at the inhibitory residues S638 and S758, a scenario that is associated with inhibition of autophagy initiation.

One aim of this study was the characterization of the role of the SIRT4-HDAC6 and SIRT4-OPA1 interactions [17, 21] in the regulation of autophagy. HDAC6, that is upregulated in SIRT4/SIRT4(H161Y) expressing cells (**Fig. 2**), functions as major regulator of autophagic and mitophagic processes [23, 45], including microtubule-based cargo/mitochondrial transport and autophagosome-lysosome fusion. Pharmacological inhibition of HDAC6 successfully restored functional autophagy in various pathophysiological systems [30–33]. However, despite greatly restored levels of acetylated α-tubulin (K40) (**Fig. S5**), tubacin-mediated inhibition of HDAC6 was unable to rescue normal LC3B-II increase and autophagic flux in SIRT4(HY161) expressing cells that were exposed to autophagy/mitophagy triggers (**Fig. 3**, **Fig. S6A and S7A**). Furthermore, the functional interaction of SIRT4 with the mitochondrially inner-membrane localized GTPase OPA1 [17, 46] also provided a potential molecular basis for the autophagy phenotype of SIRT4(H161Y) expressing cells. OPA1-L, which was clearly increased/stabilized by SIRT4(H161Y) in CoCl_2_-treated cells (**Fig. 4**), promotes mitochondrial fusion and thereby counteracts mitophagy as compared to OPA1-S that is linked to mitochondrial fission [19, 34]. However, the OPA1 inhibitor MYLS22 failed also to significantly rescue autophagy/mitophagy in our experimental systems (**Fig. 5**, **Fig. S6B and S7B**). Taken together, deregulation of neither HDAC6 nor OPA1 seems causally linked to the decreased LC3B-II levels in SIRT4(H161Y)-expressing HEK293 cells.

At first sight, our findings seem in line with a report from Liu et al., demonstrating that overexpression of SIRT4 decreases the levels of PTEN, thereby antagonizing PI3K and mTORC1 signaling and promoting autophagy [47, 48]. However, the destabilizing effect on PTEN occurs also in the case of SIRT4(H161Y) expression [47], thus being not only SIRT4 enzymatically independent, but also ruling out the SIRT4-PTEN interaction as molecular basis responsible for the autophagic phenotype in our experimental models. Of note, a direct interaction of SIRT4 with PTEN upstream of mTORC1 signaling requires a significant cytoplasmatic localization of SIRT4. The latter is actually the case, given that SIRT4 is distributed between the cytoplasm, nucleus, and mitochondria [21, 49]. Moreover, SIRT4 protein levels are controlled *via* degradation both by autophagy and the cytoplasmatic proteasome [25, 50]. Lastly, several recent reports attributed novel extramitochondrial roles to SIRT4 in microtubule dynamics and regulation of mitotic cell cycle progression [21], WNT/β-catenin and hippo signaling [51, 52], C-RAF-MAPK signaling [53], direct nuclear regulation of the mRNA splicing factor U2AF2 [54], and SNARE complex formation required for autophagosome-lysosome fusion [55].

The impact of SIRT4 on autophagy seems to be tightly linked to its role in metabolic regulation through glutamate dehydrogenase (GDH) and downstream signaling towards AMPK [56, 57]. One possibility would be that mitochondrial SIRT4 inhibits GDH through ADP-ribosylation [58], thus reducing conversion of glutamate (from glutamine) into α-ketoglutarate. As a consequence, cytosolic glutamine levels increase which in turn activate mTORC1 and inhibit autophagy [57]. Indeed, Shaw et al. described this scenario for increased SIRT4 levels under anabolic, *i.e*., high-glucose conditions [59]. The authors observed an activation of mTORC1 (due to increased cytosolic glutamine), increased inhibitory ULK1^pS758^ phosphorylation, and decreased LC3B-II, all resulting ultimately in inhibition of autophagy [59]. Another mechanism lies in the possibility, that SIRT4-mediated inhibition of GDH may result in reduced ATP levels, *i.e*., an increased AMP/ADP:ATP ratio that leads to an increased activation of AMPKα and ULK1 and thus more autophagy initiation [56]. The latter mechanism would be in line with our work and also with further findings from Wang et al. describing that the pro-autophagic SIRT4-GDH↓-AMPKα ↑-mTORC1 ↓ axis was negatively affected by expression of SIRT4(H161Y) [43]. Lastly, a recent report uncovered that SIRT4 activates autophagy also *via* AMPKα-p53 signaling [41], however, the regulation at the level of ULK1 has not been addressed. AMPK takes over a bifunctional role in ULK1 phosphorylation and autophagy regulation. On the one side, AMPK mediated phosphorylation of S777 and S317 promotes activation of ULK1 [40]. On the other side, recent work by Park et al. demonstrated that AMPK-mediated phosphorylation of UKL1 residues S556 and T660 prevents dephosphorylation of the key inhibitory residue S758 and hence ULK1 activation towards autophagy initiation [60]. Of note, the latter inhibitory mechanism of AMPK towards ULK1 has been demonstrated under glucose starvation to maintain autophagy competence and cellular survival during prolonged energy stress [60, 61].

## Conclusions

The dominant-negative mutant SIRT4(H161Y), in contrast to wild-type SIRT4, promotes inhibitory phosphorylations of ULK1 at S758 and S638 that are associated with negative regulation of autophagy initiation and subsequent LC3B-II linked autophagic flux. These findings are the basis for our working model (**Fig. 8**) in which SIRT4 positively regulates the early autophagic phase in an enzymatically dependent manner. The underlying molecular mechanism(s) of the interaction of SIRT4 with the AMPK-mTORC1-ULK1 network directly in the cytoplasm, or rather indirectly *via* mitochondrial metabolic regulation of the AMP/ATP ratio and AMPKα modulation, remain to be elucidated.

**Fig. 8.**
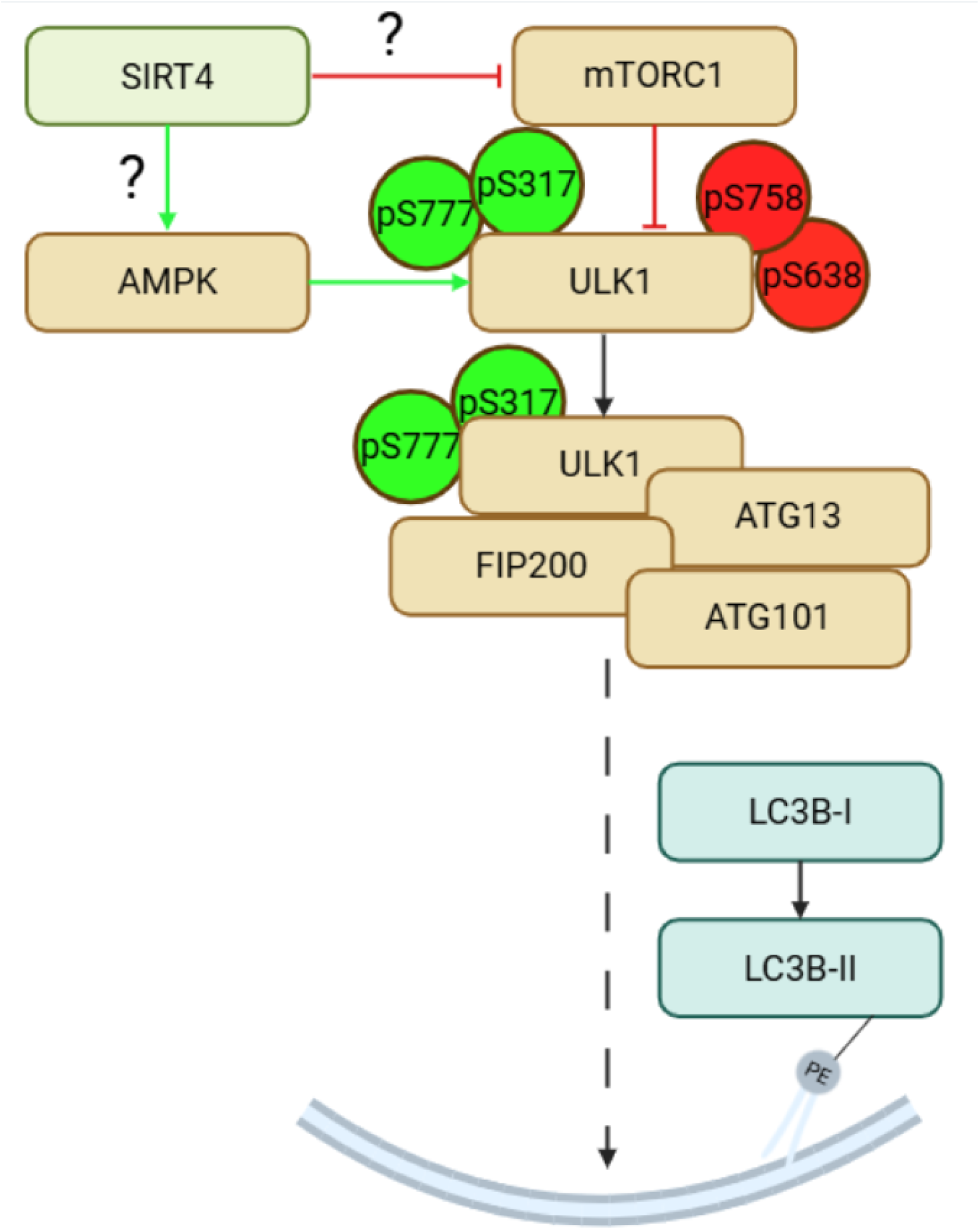
Working model suggesting a regulatory role of SIRT4 in AMPK-dependent activation and mTORC1-driven inactivation of ULK1, a central driver of autophagy initiation. Activation of ULK1 involves AMPK mediated phosphorylations of S777 and S317, whereas the mTORC1 targets S758 and S638 are linked to ULK1 inhibition. Residues S556 and T660, which can be also phosphorylated by AMPK thereby preventing dephosphorylation of the S758 and hence ULK1 activation [60], are not depicted in this model. PE, phosphatidylethanolamine anchor. Model modified after [39] and [62]. Created with BioRender.com.

## Author Contributions

R.P.P., I.L., and K.A. initiated the project and designed the study. I.L., K.A., N.H., A.I., J.T.V.N, D.M.F., and R.P.P. performed and analyzed the experiments. J.H., C.W., H.H., J.S., M.R.A, B.S., and D.M.F provided expertise and essential methods and reagents. R.P.P. wrote the manuscript. All authors read, discussed, corrected, and approved the final version of the manuscript.

## Acknowledgments

We thank Christine Cosmovici (Institute of Virology) for help with flow cytometry measurements using the mt-mKEIMA probe, and Ursula Duerkop and Yvonne Arlt for expert technical assistance. This work was funded in part by the Stiftung für Altersforschung (grant 701.810.783) of the Heinrich Heine-University Düsseldorf (to R.P.P.).

## Abbreviations

AMPK: AMP-activated protein kinase
BafA1: Bafilomycin A_1_
CCCP: carbonyl cyanide 3-chlorophenylhydrazone
GDH: Glutamate dehydrogenase
HDAC6: Histone deacetylase 6
LC3B: microtubule-associated protein 1 light chain 3 B
mTORC1: mammalian target of rapamycin complex 1
OPA1: optic atrophy 1
SIRT: sirtuin
SIRT4: sirtuin 4
ULK1: Unc-51-like kinase 1

---

**Table S1.** Primary antibodies used in this study.

| <u>Primary antibodies</u> | <u>Species</u> | <u>Dilution</u> | <u>Supplier</u> | <u>Reference</u> |
| --- | --- | --- | --- | --- |
| myc | rabbit | 1:2.000 | Cell Signaling Technology | 2272 |
| eGFP | mouse | 1:2.000 | Roche | 11814460001 |
| eGFP | mouse | 1:20.000 | Proteintech | 66002-1-AP |
| HDAC6 | rabbit | 1:1000 | Cell Signaling Technology | 7558 |
| acetyl. Tubulin (K40) | mouse | 1:1000 | Santa Cruz Biotechnology | sc-23950 |
| α-Tubulin | rabbit | 1:1000 | Abcam | ab52866 |
| α-Tubulin | rabbit | 1:2.000 | Proteintech | 11224-1-AP |
| OPA1 | mouse | 1:1000 | BD | 612607 |
| HIF1 $\alpha$ | rabbit | 1:1000 | Abcam | ab179483 |
| LC3 | rabbit | 1:500 | Cell Signaling Technology | 2775 |
| ULK1 | rabbit | 1:1000 | Cell Signaling Technology | 8054 |
| pS757-ULK | rabbit | 1:1000 | Cell Signaling Technology | 6888 |
| pS638-ULK | rabbit | 1:1000 | Cell Signaling Technology | 14205 |
| phospho-S6 (pS6 Ser240/244) ribosomal protein | rabbit | 1:1000 | Cell Signaling Technology | 2215 |
| S6 ribosomal protein | mouse | 1:1000 | Cell Signaling Technology | 2317 |
| β-Actin | mouse | 1:20.000 | Proteintech | 66009-1-Ig |

## Supplementary Figures

**Fig. S1.** CoCl_2_ treatment induces a pseudo-hypoxic response in wild-type HEK293 cells. HEK293 cells were subjected to treatment with 250 µM and 400 µM CoCl_2_ for 24 h followed by immunoblot analysis of HIF1α (**A**) and LC3B-II (**B**) protein levels.

**Fig. S2**. Generation and characterization of HEK293 cell lines stably expressing C-terminal eGFP fusion or myc-Flag tagged versions of SIRT4 or SIRT4(H161Y). (**A**) Expression levels of SIRT4-eGFP and SIRT4(H161Y)-eGFP were measured by flow cytometry (BD FACS Canto II) and results were analysed using the FlowJo v10 software and calculated as MFI (Median Fluorescence Intensity). (**B**) HEK293 cell lines stably expressing SIRT4-eGFP or SIRT4(H161Y)-eGFP were subjected to treatment with 400 µM CoCl_2_ for 24 h followed by immunoblot analysis of HIF1α protein levels. (**C**) Analysis of SIRT4-myc-Flag and SIRT4(H161Y)-myc-Flag expression levels in HEK293 cell lines by immunoblotting. (**D**) HEK293 cell lines stably expressing SIRT4-myc-Flag or SIRT4(H161Y)-myc-Flag together with the autophagic flux probe LC3-eGFP were subjected to treatment with 400 µM CoCl_2_ for 24 h followed by immunoblot analysis of HIF1α protein levels.

**Fig. S3**. Inhibitory impact of SIRT4(H161Y) on autophagic flux upon CoCl_2_-induced pseudohypoxia. (**A**) HEK293 cells stably expressing eGFP, SIRT4-eGFP, or SIRT4(H161Y)-eGFP were subjected to CoCl_2_ treatment for 36 h followed by immunoblot analysis of LC3B-I/II levels. (**B, C**) Relative quantification of immunoblot signals of LC3B-II and LC3B-I was performed using ImageJ-based densitometric evaluation and β-actin levels as loading control (n=3). To determine statistical significance, Two-Way ANOVA tests were employed (mean ± S.D.; *p < 0.05; **p < 0.01).

**Fig. S4.** SIRT4(H161Y) inhibits autophagic flux upon CCCP/oligomycin-mediated mitochondrial stress or Rapamycin treatment. HEK293 cells stably expressing myc-Flag, SIRT4-myc-Flag, or SIRT4(H161Y)-myc-Flag were subjected to CCCP/oligomycin (**A**) or rapamycin (**B**) treatment followed by flow cytometry-based analysis of autophagic flux using the GFP-LC3-RFP-LC3ΔG probe (n=7-8). To determine statistical significance, Two-Way ANOVA tests were employed (mean ± S.D.; *p < 0.05; **p < 0.01; ***p < 0.001). (**C**) HEK293 cell lines stably expressing myc-Flag, SIRT4-myc-Flag or SIRT4(H161Y)-myc-Flag were subjected to treatment with 1.25 µM rapamycin for 24 h followed by immunoblot analysis of pS6 (Ser240/244) and total ribosomal S6 protein levels.

**Fig. S5.** Immunoblot analysis of acetylated α-Tubulin (K40) protein levels in CoCl_2_-treated HEK293 cell lines expressing eGFP-fused (**A**) or myc-Flag-tagged (**B**) SIRT4 or SIRT4(H161Y) in the presence or absence of the HDAC6 inhibitor tubacin. Probing against α-tubulin was used as loading control.

**Fig. S6.** Impact of tubacin or MYLS22 treatment on the inhibited autophagic flux of CCCP/oligomycin treated HEK293-SIRT4(H161Y) cells. HEK293 cells stably expressing myc-Flag, SIRT4-myc-Flag, or SIRT4(H161Y)-myc-Flag were subjected to CCCP/oligomycin treatment in the presence of tubacin (HDAC6 inhibitor; n=6) (**A**) or MYLS22 (OPA1 inhibitor; n=4) (**B**) followed by flow cytometry-based analysis of autophagic flux using the GFP-LC3-RFP-LC3ΔG probe. To determine statistical significance, Two-Way ANOVA tests were employed (mean ± S.D.; *p < 0.05; *** p < 0.001).

**Fig. S7.** Impact of tubacin or MYLS22 treatment on the inhibited autophagic flux of rapamycin treated HEK293-SIRT4(H161Y) cells. HEK293 cells stably expressing myc-Flag, SIRT4-myc-Flag, or SIRT4(H161Y)-myc-Flag were subjected to rapamycin treatment in the presence of tubacin (HDAC6 inhibitor; n=6) (**A**) or MYLS22 (OPA1 inhibitor; n=4) (**B**) followed by flow cytometry-based analysis of autophagic flux using the GFP-LC3-RFP-LC3ΔG probe. To determine statistical significance, Two-Way ANOVA tests were employed (mean ± S.D.; *p < 0.05; **p < 0.01; ***p < 0.001).

