## Supplementary figures and images for "SIRT4 positively regulates autophagy *via* ULK1, but independently of HDAC6 and OPA1"

### Supplemental Figures

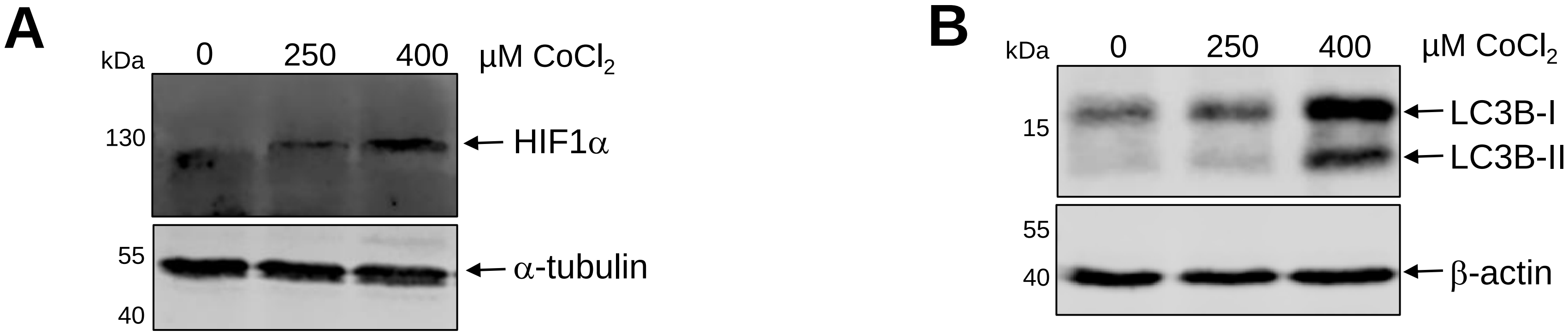

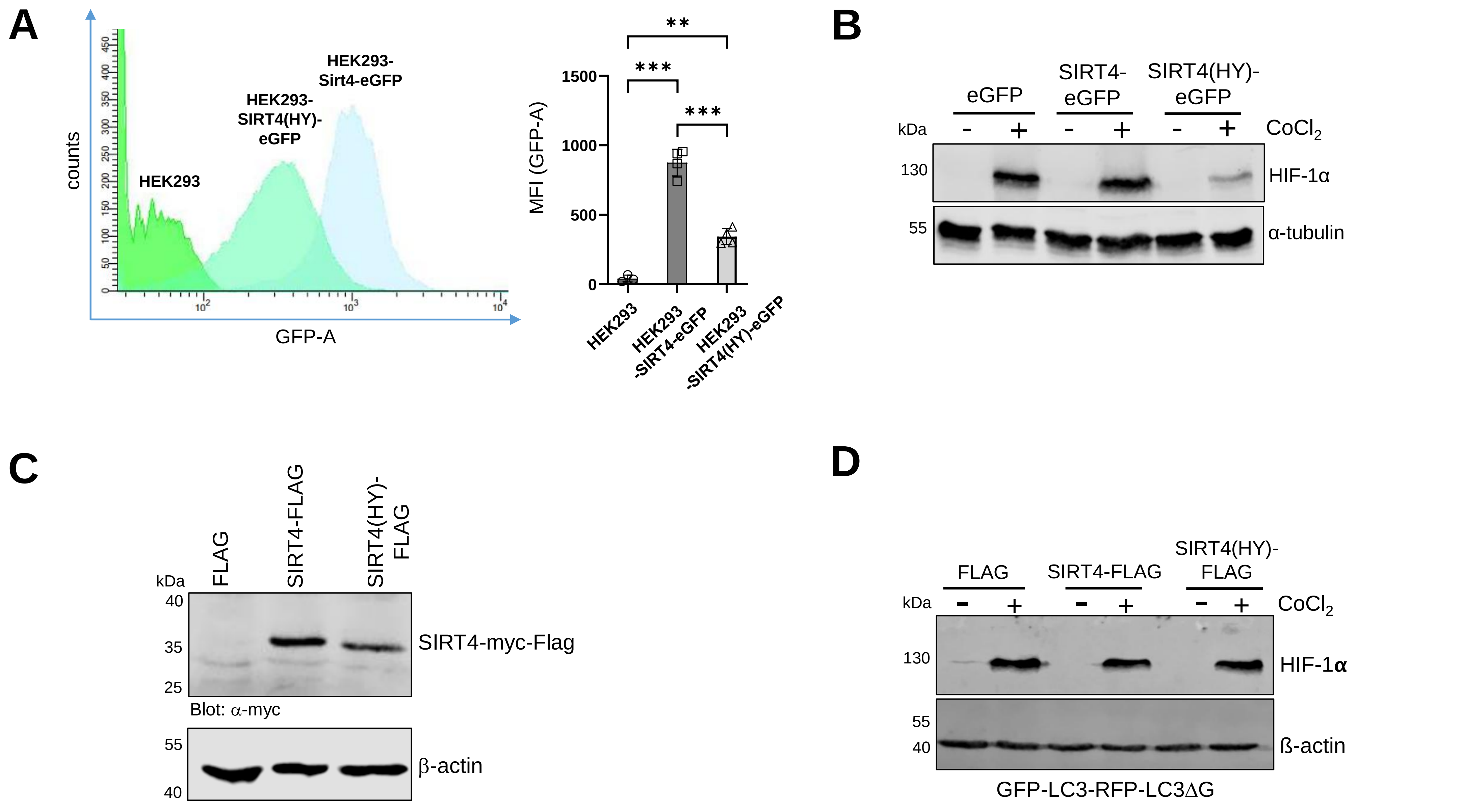

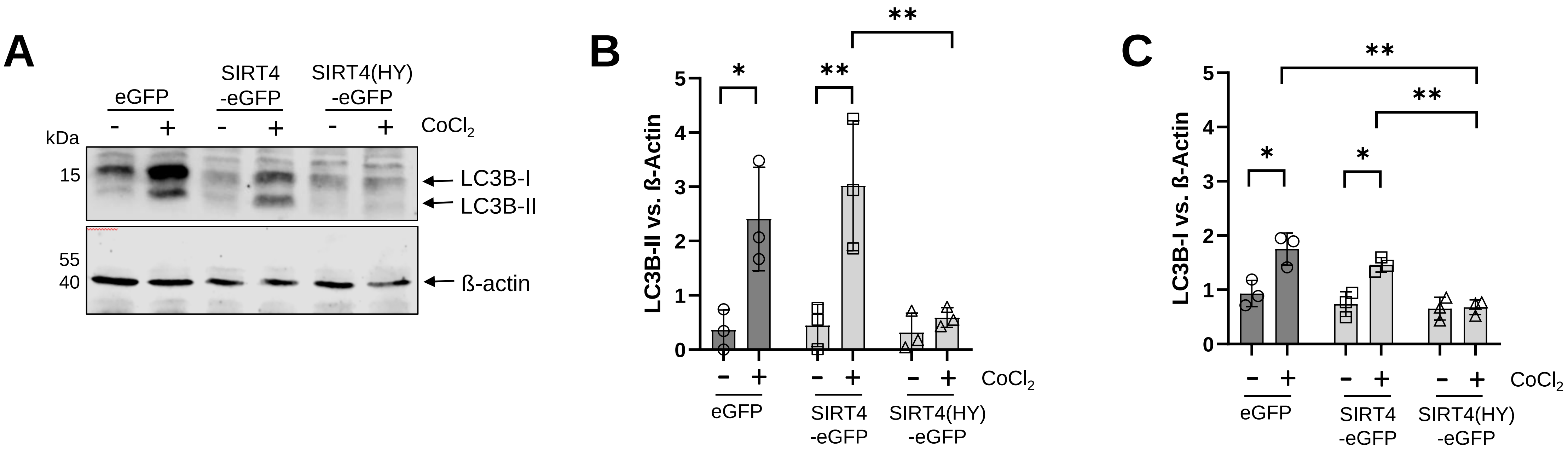

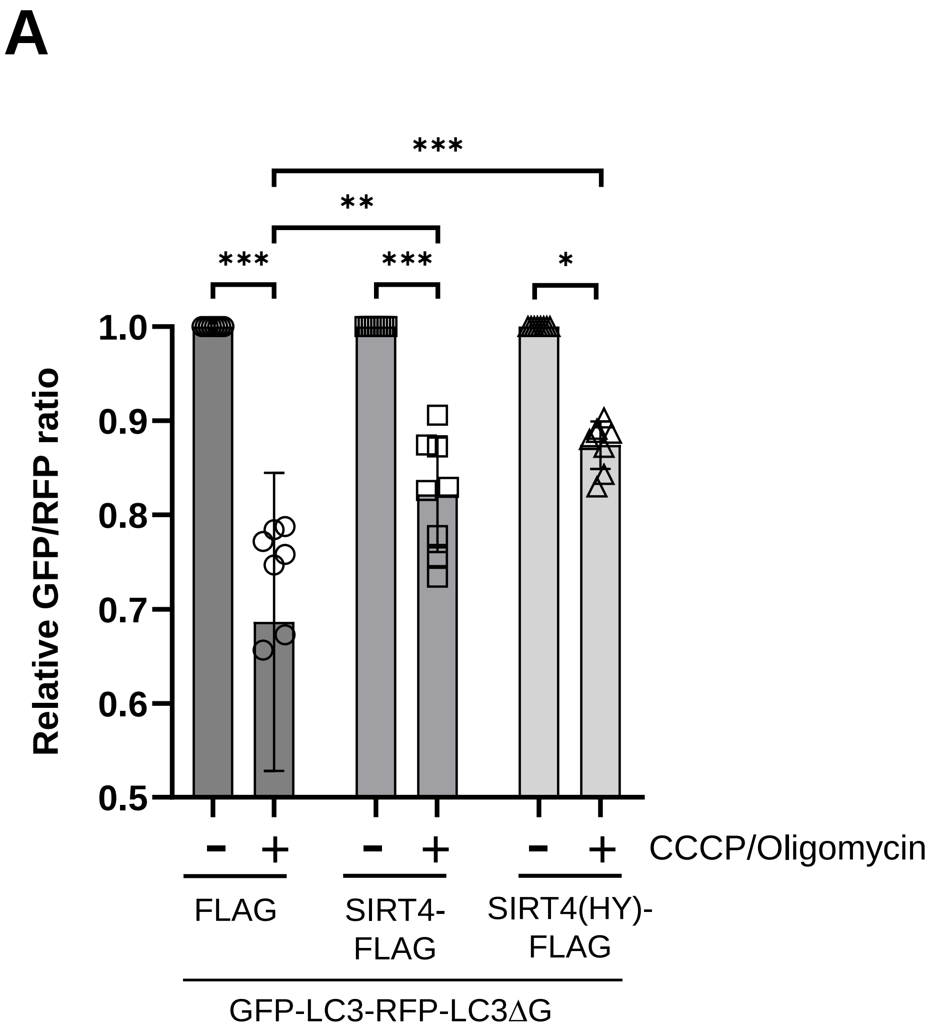

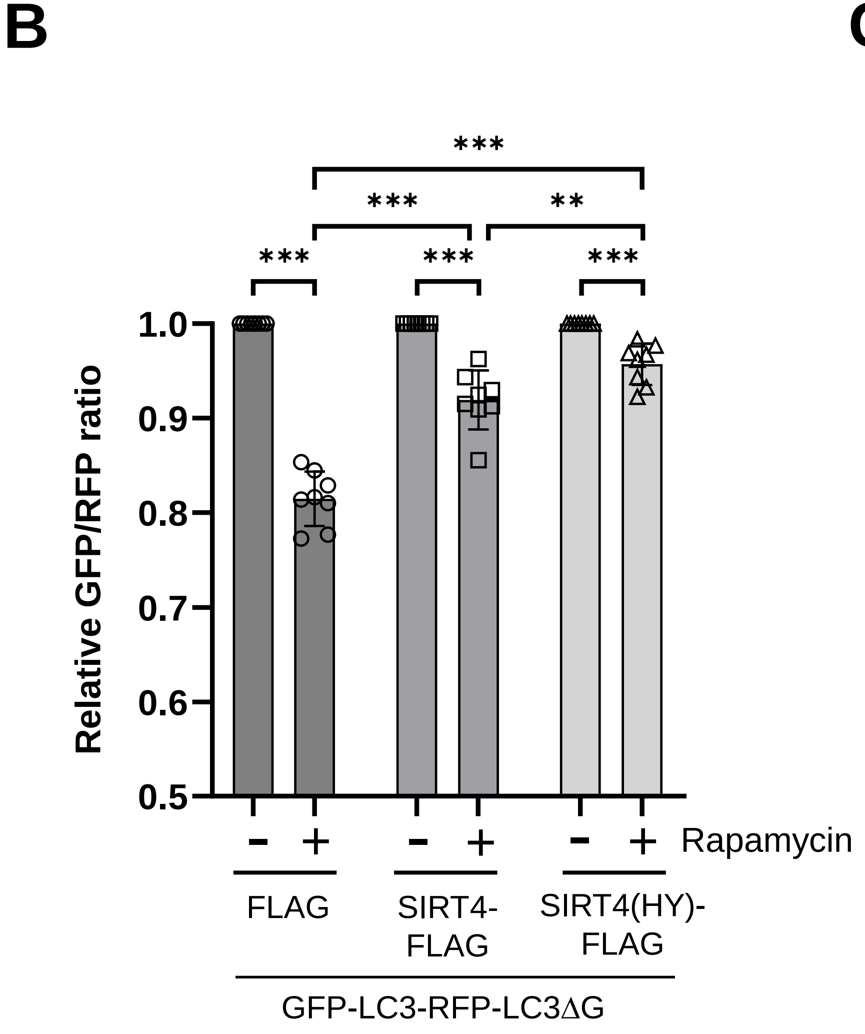

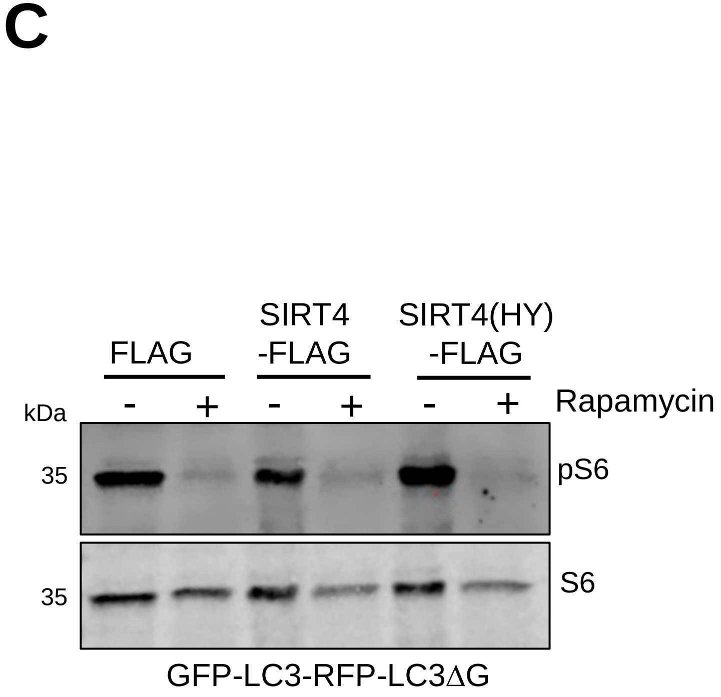

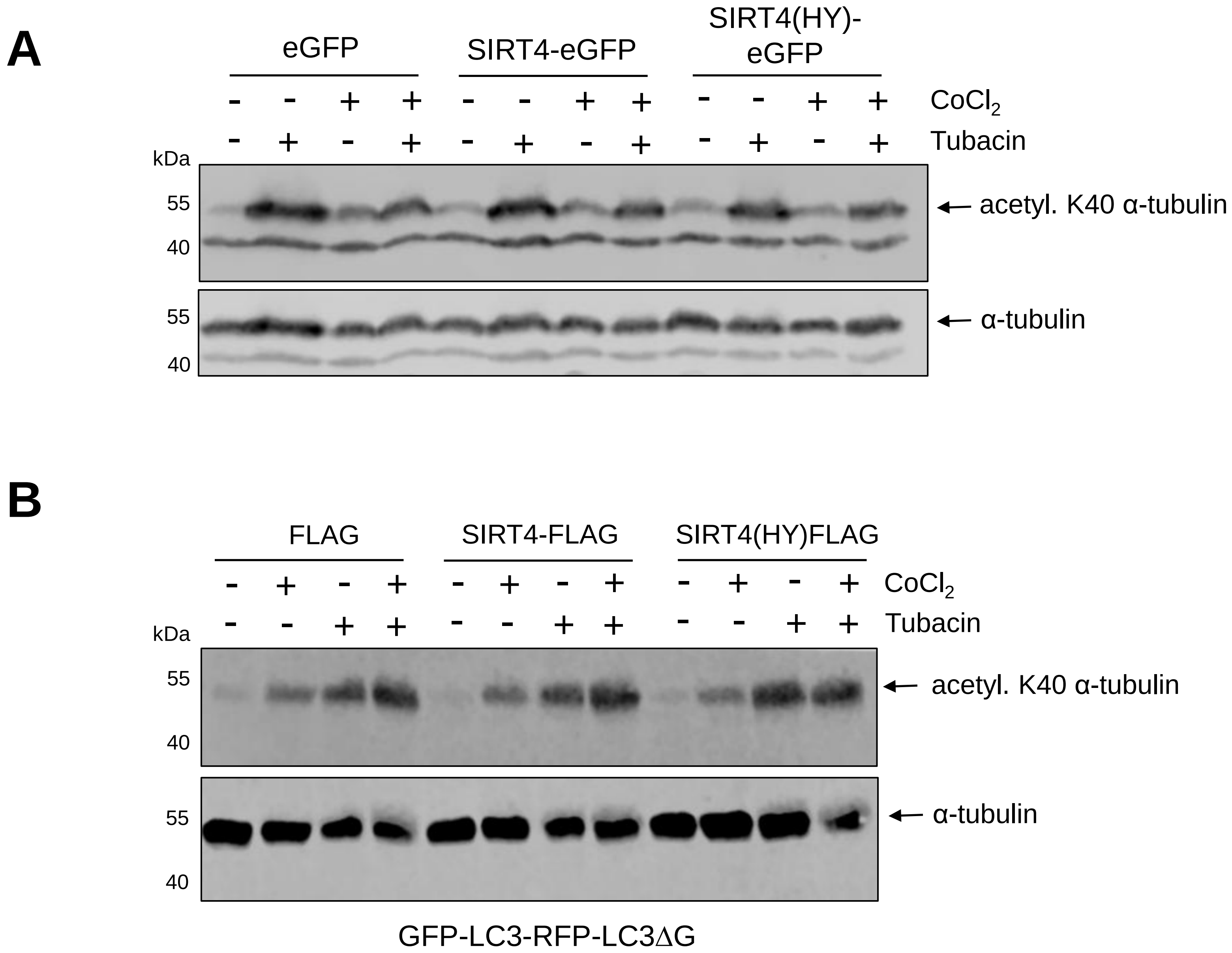

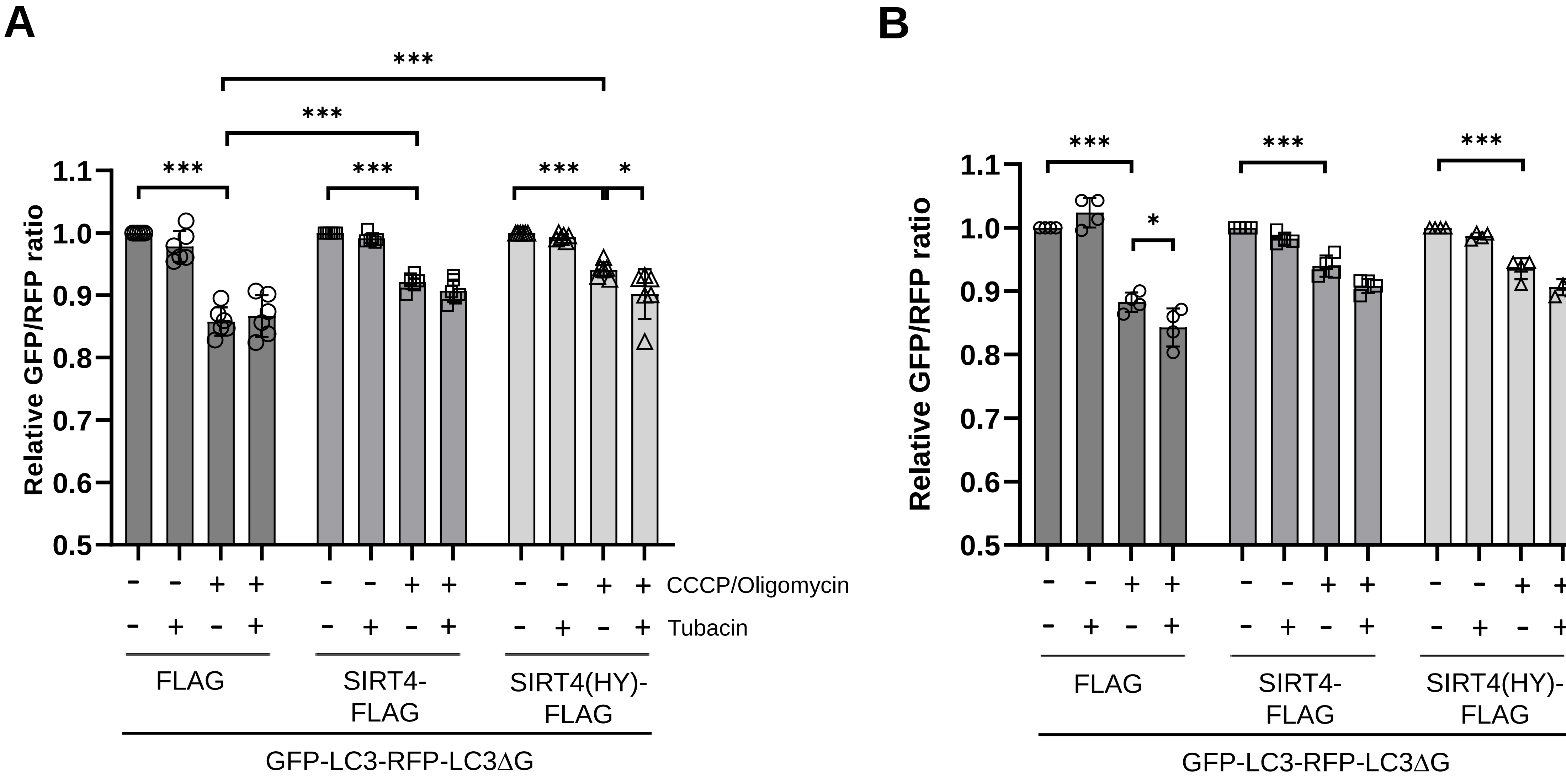

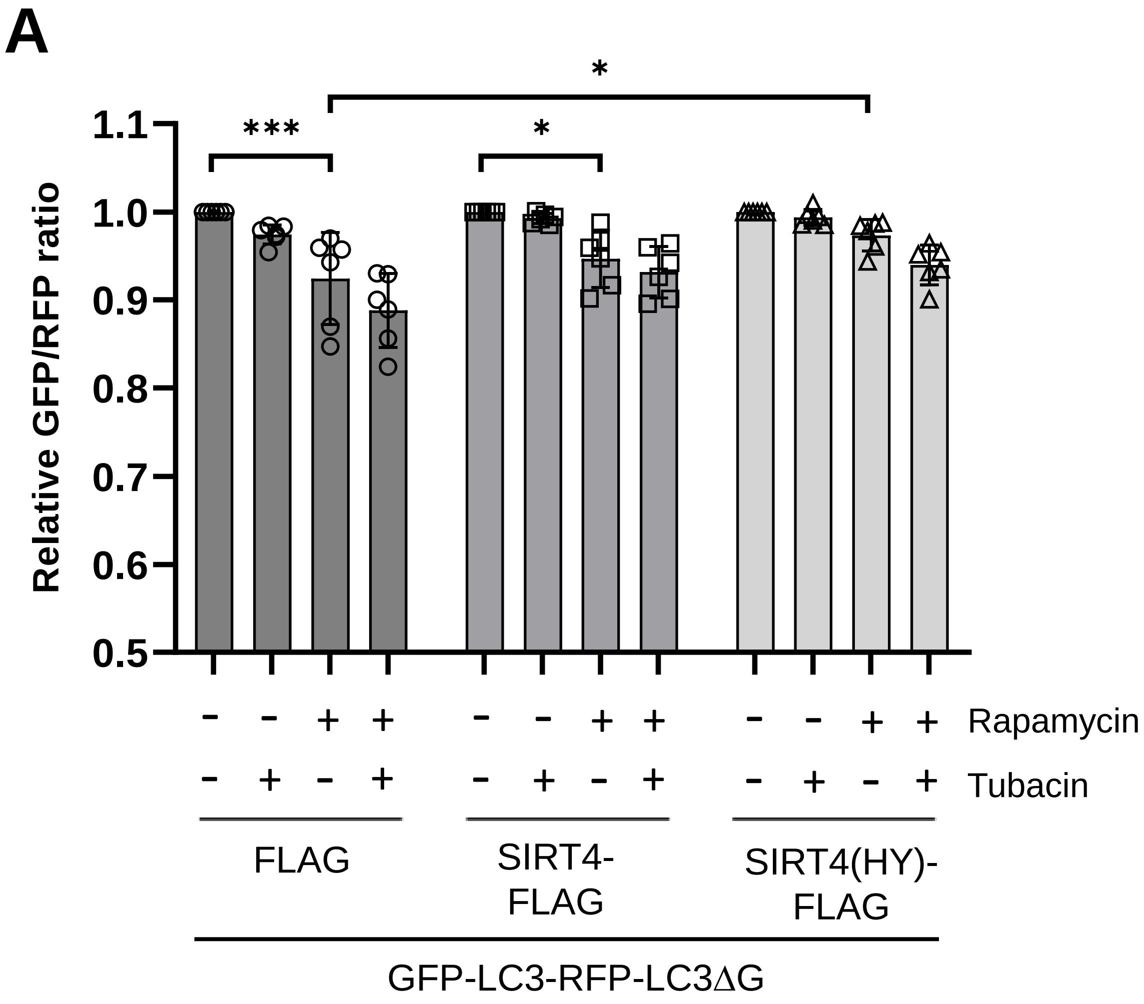

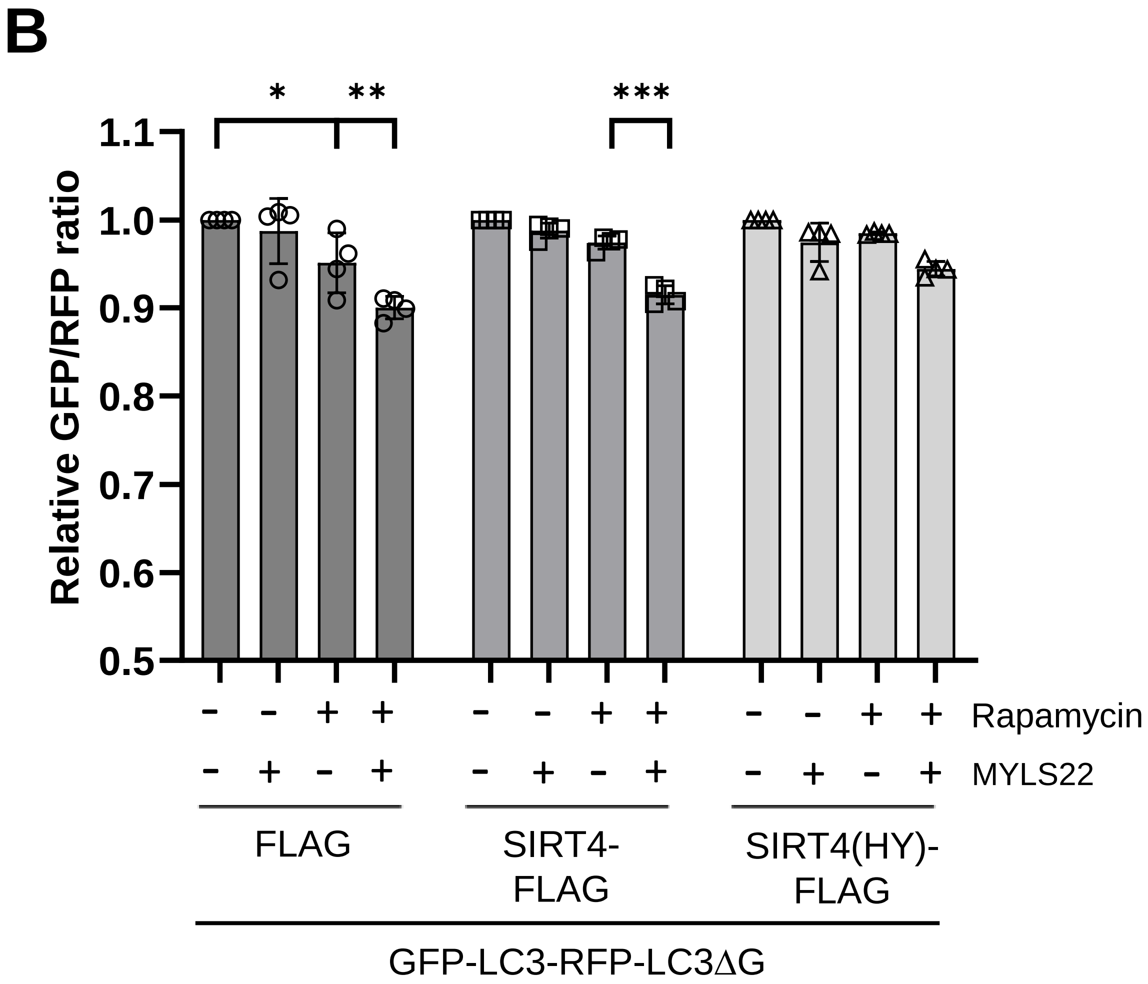
